# Functional plasticity of HCO_3_^-^ uptake and CO_2_ fixation in *Cupriavidus necator* H16

**DOI:** 10.1101/2024.05.07.593039

**Authors:** Justin Panich, Emili Toppari, Sara Tejedor-Sanz, Bonnie Fong, Eli Dugan, Yan Chen, Christopher J. Petzold, Zhiying Zhao, Yasuo Yoshikuni, David F. Savage, Steven W. Singer

## Abstract

Uptake and fixation of CO_2_ are central to strategies for CO_2_-based biomanufacturing. *Cupriavidus necator* H16 has emerged as a promising industrial host for this purpose. Despite its prominence, the ability to engineer *C. necator* inorganic carbon uptake and fixation is underexplored. Here, we test the role of endogenous and heterologous genes on *C. necator* inorganic carbon metabolism. Deletion of one of the four carbonic anhydrases in *C. necator*, β-carbonic anhydrase *can*, had the most deleterious effect on *C. necator* autotrophic growth. Replacement of this native uptake system with several classes of dissolved inorganic carbon (DIC) transporters from *Cyanobacteria* and chemolithoautotrophic bacteria recovered autotrophic growth and supported higher cell densities compared to wild-type (WT) *C. necator* in saturating CO_2_ in batch culture. Several heterologous strains with *Halothiobacillus neopolitanus* DAB2 (hnDAB2) expressed from the chromosome in combination with diverse rubisco homologs grew in CO_2_ equally or better than the wild-type strain. Our experiments suggest that the primary role of Can carbonic anhydrase during autotrophic growth is for bicarbonate accumulation to support anaplerotic metabolism, and an array of DIC transporters can complement this function. This work demonstrates flexibility in HCO_3_^-^ uptake and CO_2_ fixation in *C. necator*, providing new pathways for CO_2_-based biomanufacturing.

**Graphical abstract:** 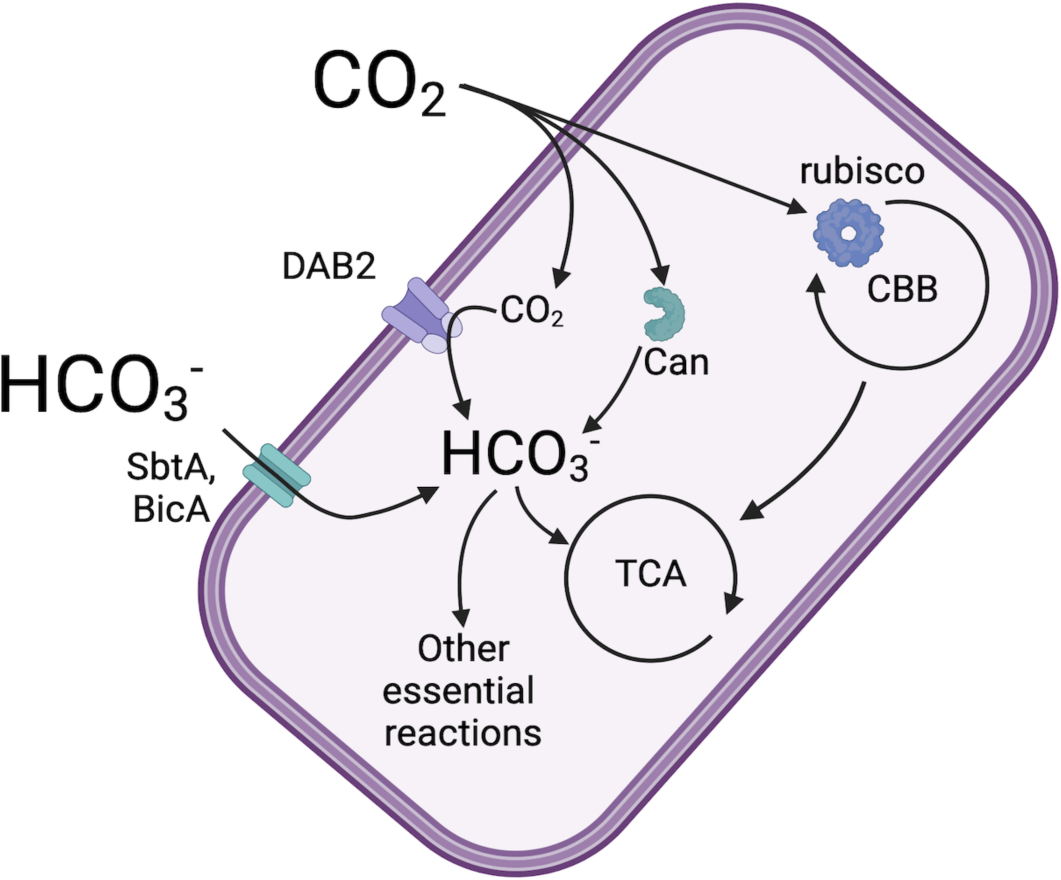

## Introduction

Society relies on chemical processes that emit vast quantities of CO_2_ and other greenhouse gases, contributing substantially to climate change^1,2^. In 2022, roughly 36.8 billion tons of CO_2_ were released into the atmosphere from transportation and heavy industry^3,4^. Rapid innovation is needed to establish sustainable commodity chemical production at an industrial scale to mitigate climate change caused by greenhouse gas emissions. An attractive strategy is to use gas feedstocks, such as CO_2_, CO, CH_4_, and H_2_, for the biological synthesis of economically crucial chemicals, such as fuels and platform chemicals. These feedstocks offer several advantages: abundance, potential capture from industrial point sources (e.g., ethanol plants, cement factories, steel mills), and generation through electrolysis. The realization of a “C1- based” commodity chemical industry, reliant on one-carbon feedstocks, would mitigate climate change while stimulating economic activity.

*Cupriavidus necator* H16 (formerly *Ralstonia eutropha* H16) is a well-characterized model organism for C1-based biological conversion due to its ability to grow on CO_2_/H_2_ or formate as sole carbon and energy sources^56,7^. CO_2_ enters the cell via passive diffusion, where it is converted to dissolved inorganic carbon (DIC; HCO_3_^-^, H_2_CO_3_) through the action of cytosolic carbonic anhydrases (CAs). These metalloenzymes interconvert CO_2_ and soluble HCO_3_^-^, and are essential in most organisms because HCO_3_^-^ is a cofactor for several reactions in the tricarboxylic acid cycle (e.g. phosphoenolpyruvate carboxylase, carbamoyl phosphate synthetase, 5-amino-imidazole ribotide carboxylase, and biotin carboxylase)^8–11^. While CO_2_ spontaneously hydrates to HCO_3_^-^ at physiological pH, the HCO_3_^-^ flux requirements are not satisfied by uncatalyzed CO_2_ hydration.

CO_2_ can also be fixed by *C. necator* through the action of ribulose-1,5-bisphosphate carboxylase/oxygenase (rubisco) to generate biomass using the Calvin-Benson-Bassham (CBB) cycle. Rubisco is a relatively slow enzyme (*k_cat_* values for CO_2_ are usually 1-22 s^-1^) and catalyzes an off-target reaction with oxygen, producing a toxic intermediate 2-phosphoglycolate, which must be salvaged^12–14^. Carbon fixed from the CBB cycle can be diverted to create sustainable bioproducts using metabolic engineering. *C. necator* has recently been engineered to produce 1,3-butanediol, trehalose, 3-hydroxyproponate, myo-inositol, sucrose, glucose, sesquiterpenes, modified polyhydroxyalkanoates, and lipochitooligosaccharides using CO_2_ feedstocks^15–22^. While this organism has proven useful for CO_2_ bioconversion, it has never been fully optimized and typically yields low titers of target molecules (often <1 g/L) while requiring >5% CO_2_^14,23^.

*C. necator* encodes four carbonic anhydrases (*caa*, *can, can2, cag*) whose function and relevance to autotrophy remain poorly characterized. *C. necator* also has a cytosolic rubisco with a relatively high specificity for CO_2_ (*S_c/o_*=75), albeit with a slower rate of catalysis compared to typical cyanobacterial rubiscos (*k_cat_*=3.8 s^-1^)^24–27^. In contrast, *Cyanobacteria* and some chemolithotrophic bacteria have evolved sophisticated CO_2_ acquisition and utilization mechanisms, known as “CO_2_ Concentrating Mechanisms” (CCMs), enabling robust growth in ambient CO_2_^28–30^. Bacteria with CCMs acquire CO_2_ in the form of HCO_3_^-^ using DIC transporters, a group of evolutionarily diverse membrane transporters that accumulate HCO_3_^-^ in the cytoplasm. HCO_3_^-^ then diffuses into a proteinaceous bacterial microcompartment, the carboxysome, where internalized CAs convert the HCO_3_^-^ to CO_2_ near the rubisco active site. The high local concentration of CO_2_ facilitates on-target rubisco carboxylation, diminishes oxygenation activity, and avoids the accumulation of the toxic intermediate 2-phosphoglycolate.

Five types of DIC transporters have been characterized to date. BCT1-like complexes use ATP to energize bicarbonate transport^31^. Other DIC transporters are Na^+^/HCO_3_^-^ symporters, such as SbtA and BicA^32,33^. CUP transporters are thylakoid membrane-associated multi-subunit complexes thought to convert H_2_O + CO_2_ into HCO_3_^-^ and a proton in alkaline conditions^34^. The fifth class of DIC transporter, the DAB complex (<u>D</u>ABs <u>A</u>ccumulate <u>B</u>icarbonate), is a heterodimeric protein thought to convert CO_2_ to HCO_3_^-^ on the cytoplasmic face of the cellular membrane using a proton gradient. DABs have been recently shown to function as the predominant DIC accumulation mechanism in *H. neopolitanus*^35^. To date, DABs have been identified in 17 phyla in *Bacteria* and *Archaea* using bioinformatic analysis and have been described for diverse organisms such as *Staphylococcus aureus, Hydrogenovibrio crunogenus,* and in heterotrophic pathogens such as *Bacillus anthracis* and *Vibrio cholera*^35–38^.

We have previously demonstrated that the robustness of autotrophic growth in three different microbes is co-limited by CO_2_ and HCO_3_^-^ availability^39,40^. In the current study, we expand upon these findings by elucidating the individual contributions of each of the four carbonic anhydrases in *C. necator*. We show that many DIC transporters can effectively replace CAs during autotrophic metabolism. We conclude that a significant limitation for autotrophic growth in *C. necator* is HCO_3_^-^ availability.

## Results

### Strains and plasmids

**Table 1.**
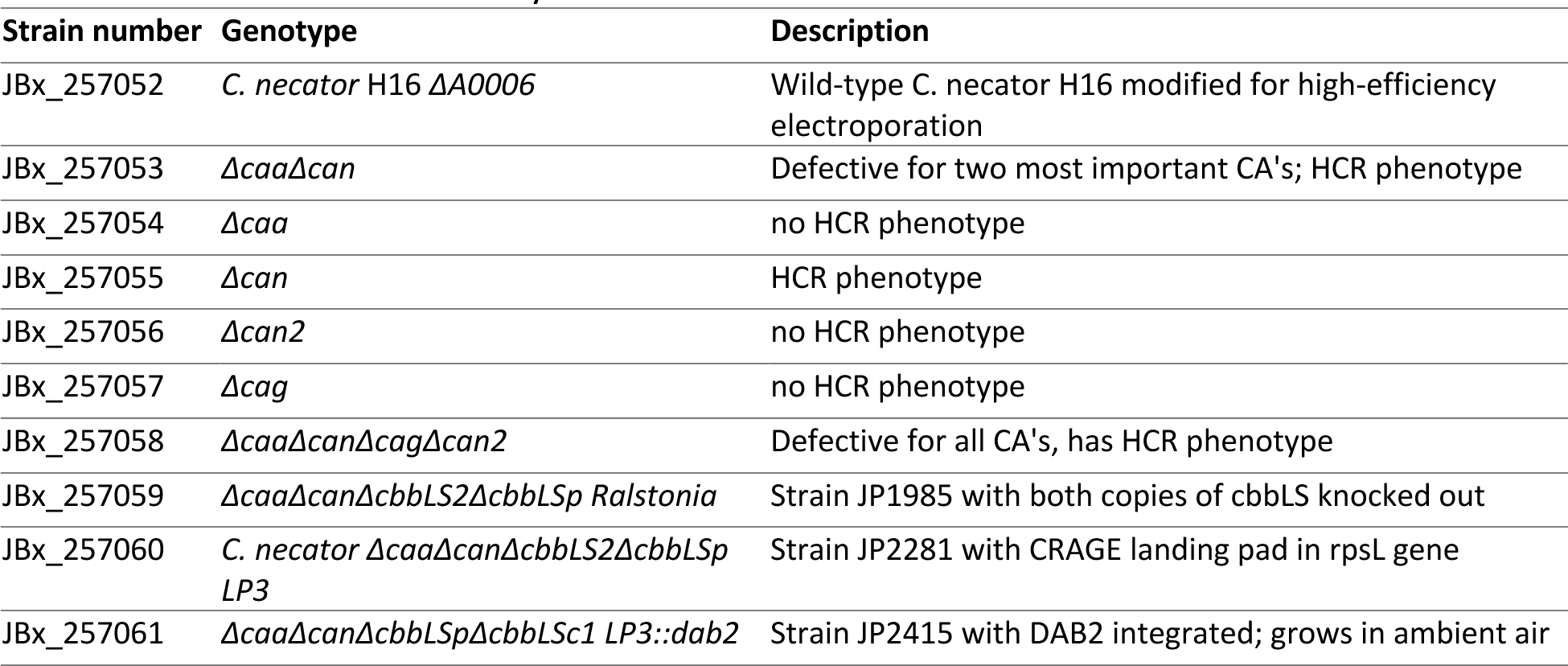
Strains used in this study.

**Table 2.**
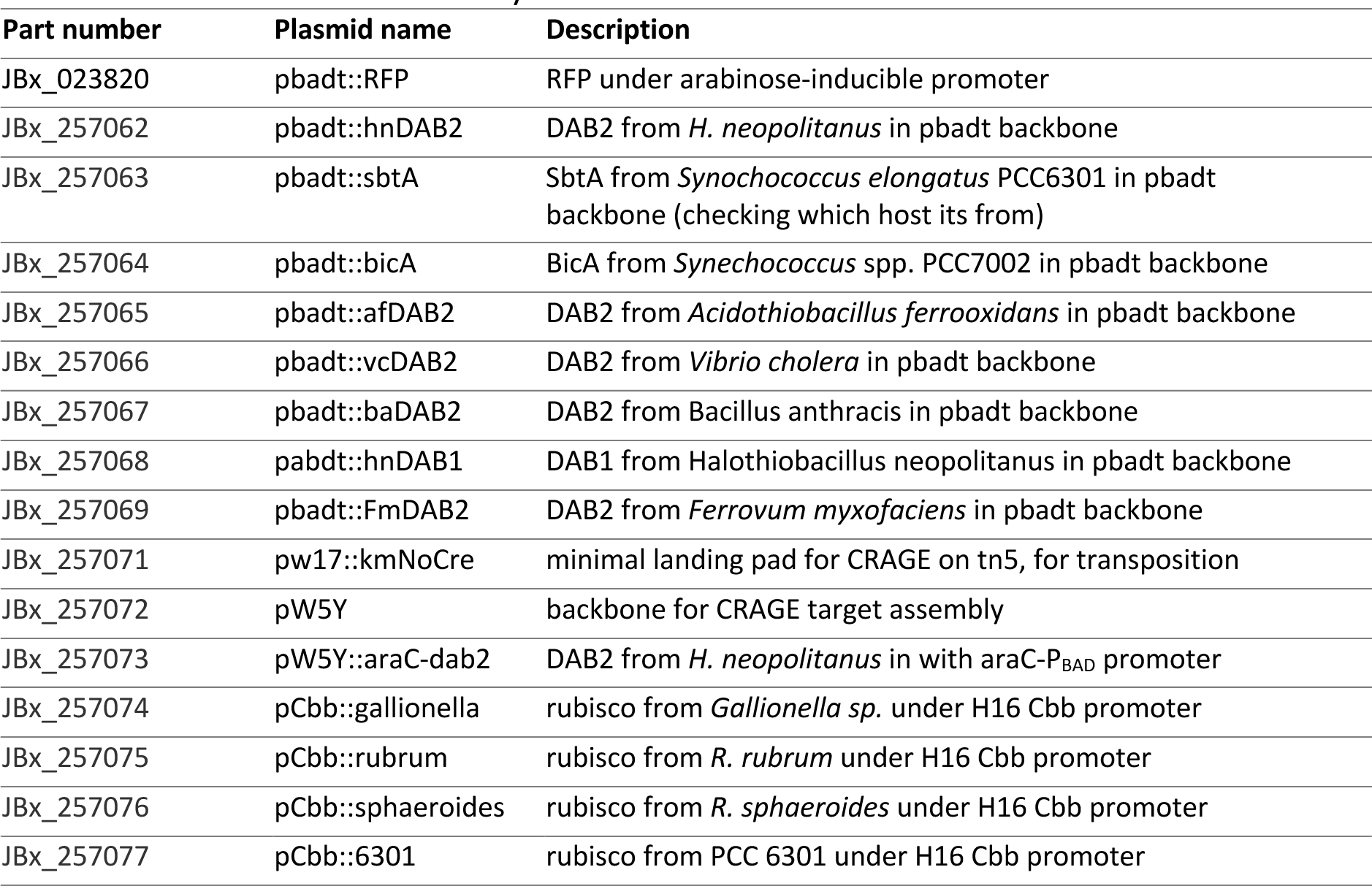
Plasmids used in this study.

### Can and Caa are the most relevant CAs for autotrophic growth in C. necator

Prior studies have interrogated the function of each of the four carbonic anhydrases found in the *C. necator* genome during heterotrophic and autotrophic growth^11,24^. In agreement with previously published studies, a high CO_2_ requiring (HCR) phenotype occurred during routine cultivation when *can* was deleted. Individual deletions of *caa, can2,* or *cag* did not significantly affect heterotrophic growth in ambient air. Deletion of all four carbonic anhydrases (*ΔcaaΔcanΔcagΔcan2*) had the same phenotypic effect as the *Δcan* mutant.

We next examined the growth of individual knockouts and the quadruple knockout in autotrophic growth conditions in increasing CO_2_ partial pressures (0.05%, 0.5%, 1.5%, and 5% CO_2_) by measuring the OD_600_ of each culture after 48 hours of autotrophic growth in sealed flasks, with cultures inoculated at an OD_600_=0.10 from heterotrophic precultures. Growth was poor for all strains at 0.05% CO_2_, likely because the cells consume the CO_2_ in the sealed flasks before appreciable growth occurs. The *Δcan2* and *Δcag* strains grew to a similar endpoint as the WT strain. The *Δcan* strain showed an apparent growth defect at 0.5% CO_2_ and higher and also exhibited decreased turbidity at 0.05% CO_2_ after 48 hours of incubation. Similarly, deletion of *caa* caused only a minor growth defect but showed robust growth at 1.5% CO_2_ (**Figure 1A**). To confirm the role of Can for autotrophic growth, we complemented the *ΔcaaΔcan* and *ΔcaaΔcanΔcan2Δcag* strains with a Can plasmid and found that the expression of Can is sufficient to support autotrophic growth in both strains (**Figure 1B**).

**Figure 1A.**
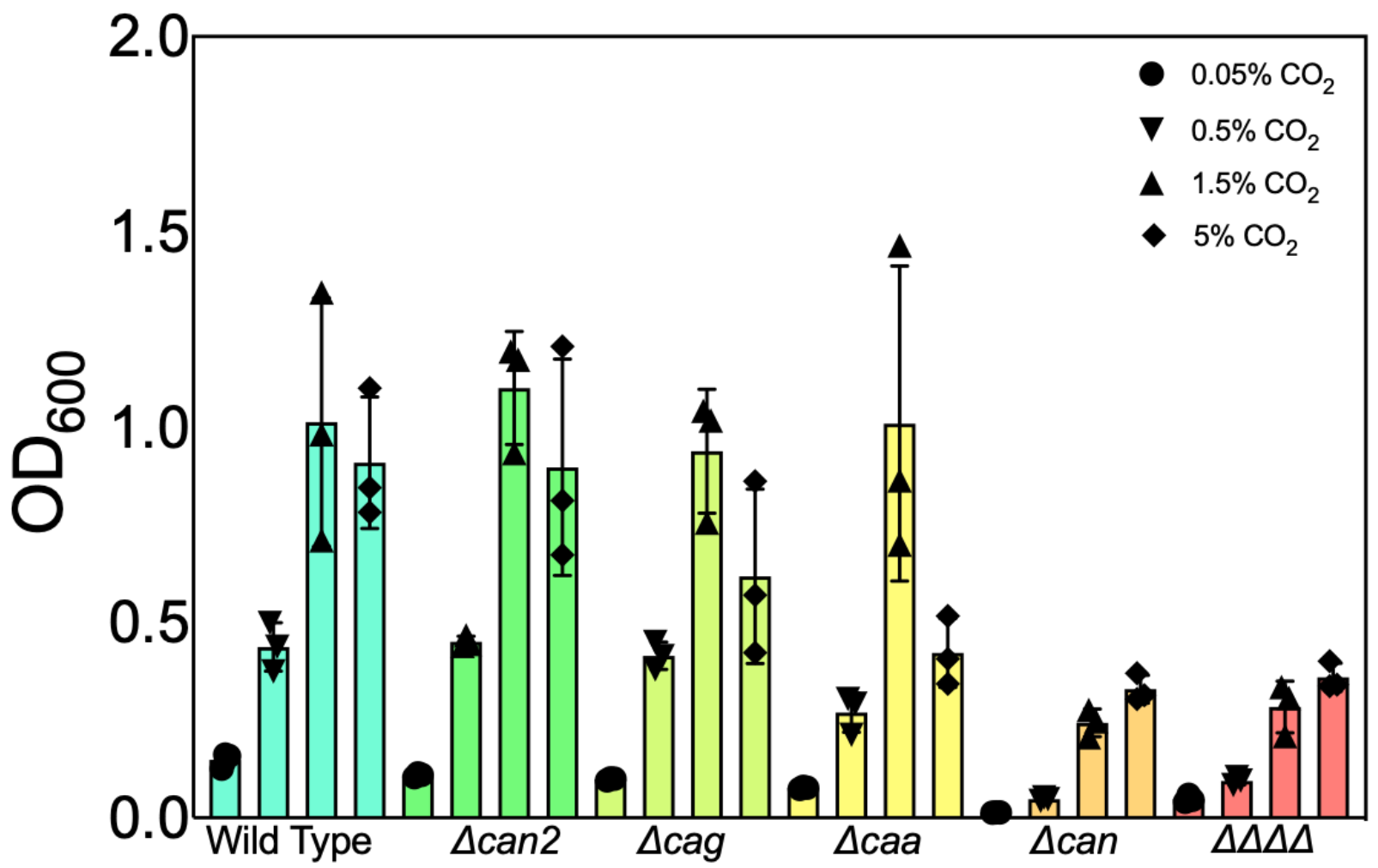
Growth profiles of CA deletion mutants. Terminal OD_600_ at 48 hours is shown in 62% H_2_/10% O_2_ and increasing CO_2_ partial pressure. Cells were inoculated from heterotrophic cultures at OD_600_ = 0.1.

**Figure 1B.**
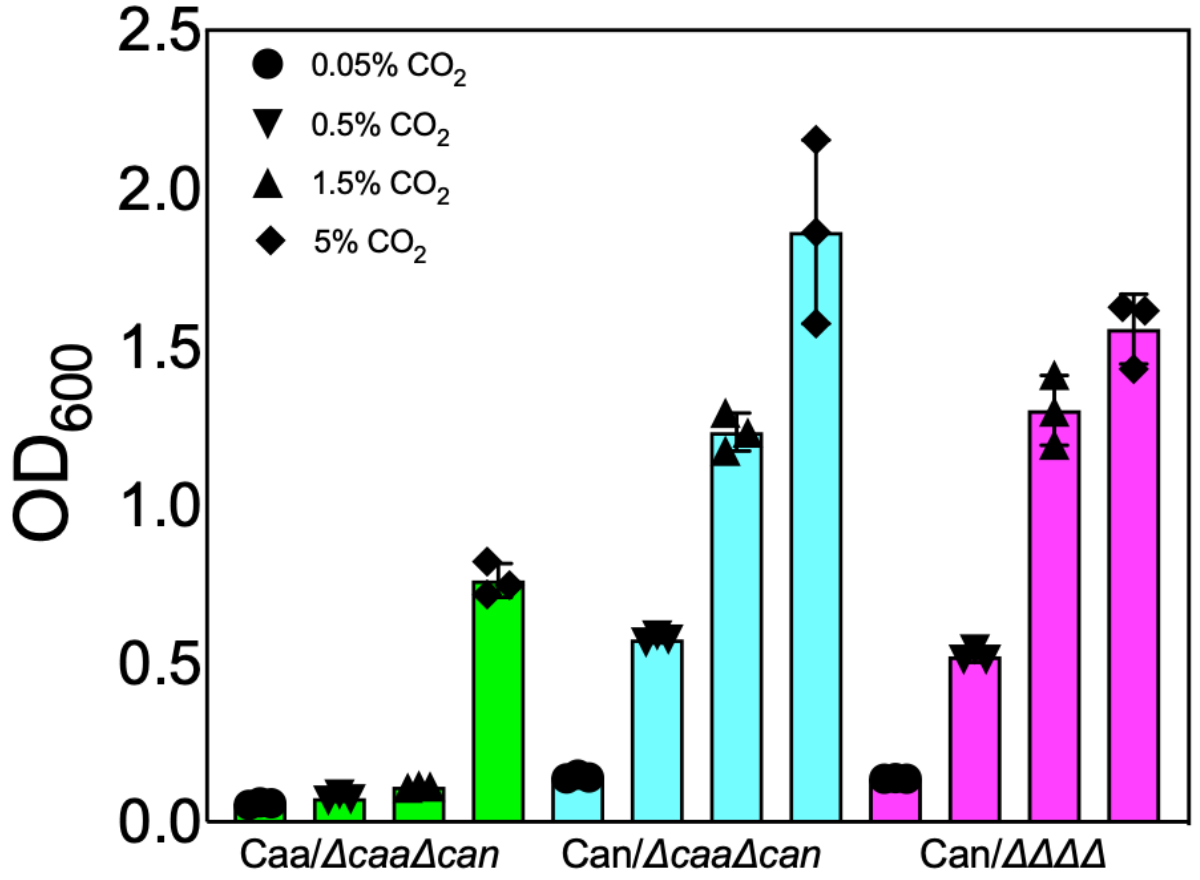
Can is sufficient for autotrophic growth, while Caa is not sufficient or necessary. Can and Caa were cloned on a pBBR1-MCS-based vector and were introduced into either *ΔcaaΔcan* or the quadruple CA deletion mutant (ΔΔΔΔ). Cells were inoculated at OD_600_ = 0.1 from heterotrophic cultures, and terminal OD_600_ values were measured after 48 hr.

### A suite of dissolved inorganic carbon transporters promotes C. necator autotrophic growth

To assess the functionality of each DIC transporter in *C. necator*, we chose a strain lacking the two most essential CAs for autotrophic growth (*ΔcanΔcaa*). We assessed the performance of six DAB-type transporters, a SbtA homolog from *Synechococcus elongatus* PCC 6301, and a BicA homolog from *Synechococcus* sp. PCC 7002^41,42^. The expression of DAB2 from *H. neopolitanus* (hnDAB2) showed comparable growth to WT cells at lower CO_2_ partial pressures (0.05% and 5% CO_2_). However, growth surpassed WT in CO_2_ in high partial pressures of CO_2_ (1.5% and 5%) (**Figure 2A**). Proteome analysis does not indicate that the expression of the two remaining CAs was impacted by hnDAB2 expression (**Figure 2B-2D**). We also found that hnDAB2 could complement the quadruple knockout strain, eliminating any substantial role for other known CAs in DIC transporter-enabled autotrophic growth (Figure S2). Proteomics revealed that rubisco expression is suppressed in 0.05% CO_2_, and 0.5% CO_2_ in the DAB2/*ΔcaaΔcan* strain relative to WT, but this data is difficult to interpret because recent work has shown that expression of rubisco is not a limiting factor in *C. necator* when grown autotrophically^43^.

**Figure 2.**
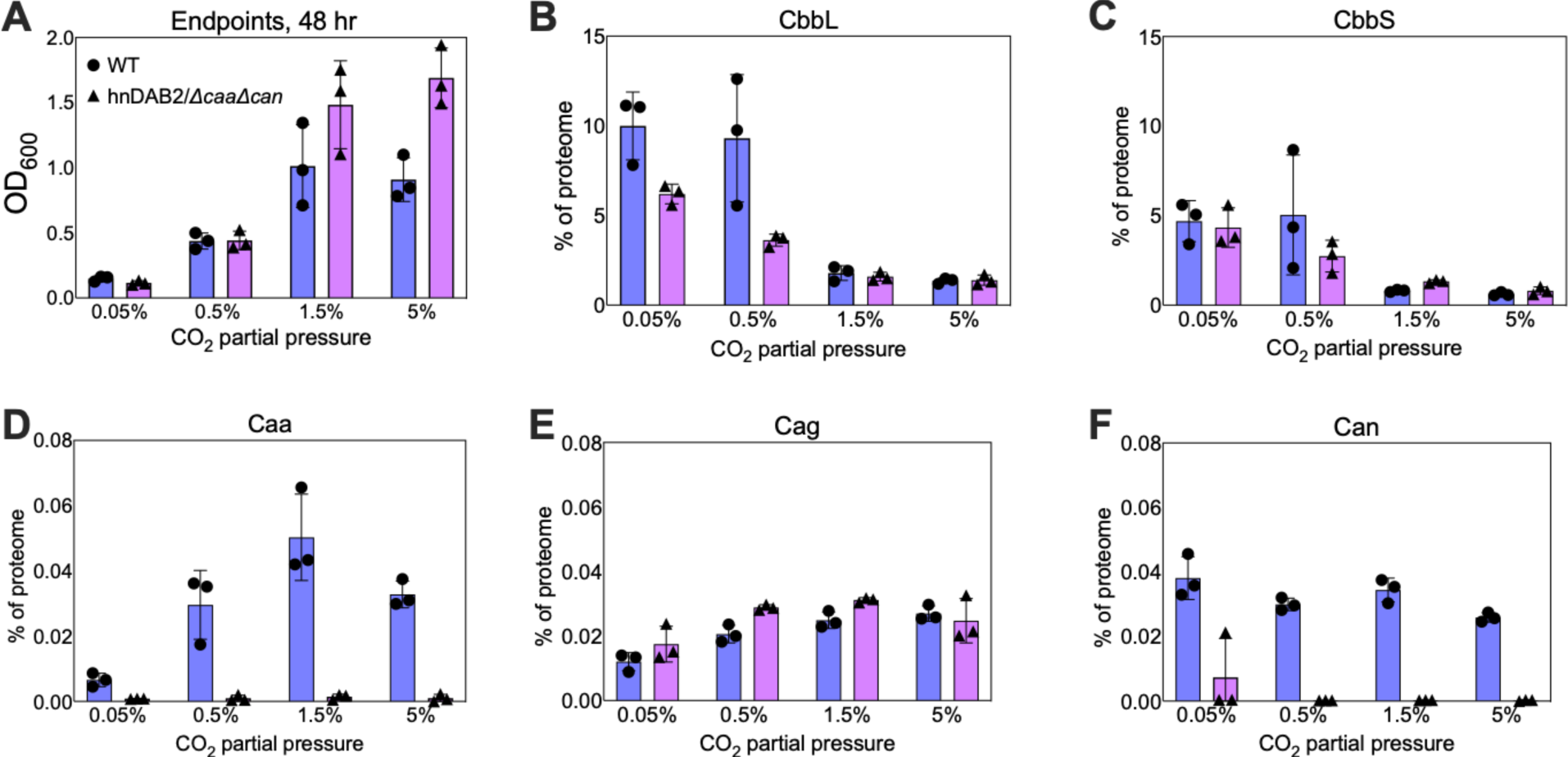
Comparison of growth characteristics and expression of key enzymes for autotrophy in wild-type and hnDAB2-complemented CA deficient mutant (*ΔcaaΔcan*). Panel A) Summary of growth data from previous figures of terminal endpoints at 48 hr in autotrophic cultures in WT or hnDAB2/*ΔcaaΔcan*. **Panel B, C)** Expression profiles for the large subunit (CbbL) and small subunit (CbbS) of rubisco **Panel D)** Expression of Caa seems to be influenced by CO_2_ partial pressure in WT cells but is knocked out in the hnDAB2-expressing strain. **Panel E, F)** Cag and Can expression do not seem to be affected by CO_2_ partial pressure or expression of hnDAB2. Can2 was not detected in these experiments.

The performance of many DIC transporters was similar: hnDAB2, atDAB2, SbtA, and BicA resulted in endpoint OD_600_ values surpassing WT *C. necator* at 1.5% and 5% CO_2_ (**Figure 3**). The DAB2 homolog from *Ferrovum myxofaciens* also exceeded the growth of WT in 5% CO_2_ but performed worse than WT in 0.05%-1.5% CO_2_. In contrast, expression of DAB2 homologs from heterotrophic pathogens *Bacillus anthracis* and *Vibrio cholera* in the *ΔcaaΔcan* background gave poor growth, despite a previous report of high activity for these homologs in a CA-free strain of *E. coli*^35^. Similarly, DAB1 from *H. neapolitanus* failed to rescue the growth of *ΔcaaΔcan*.

**Figure 3.**
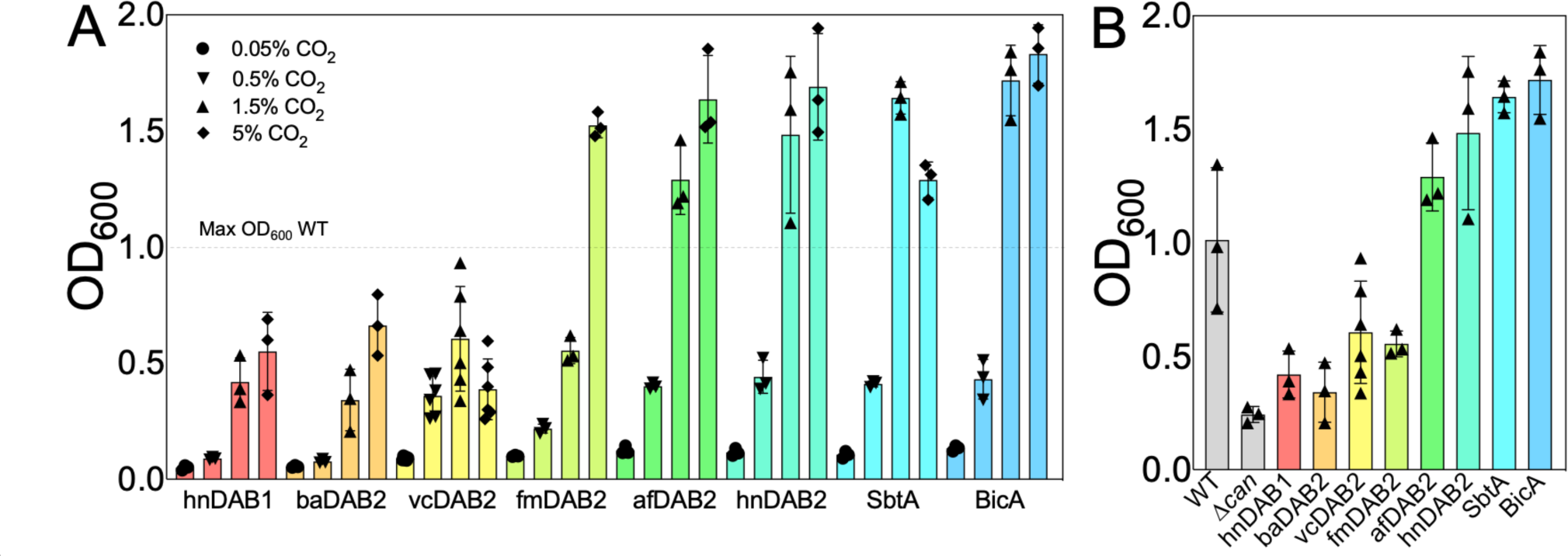
Performance of heterologous bicarbonate transporters in *ΔcaaΔcan* (ΔCA). Panel A) Autotrophic endpoints in batch culture were measured after 48 hr of autotrophic growth in 62% H_2_/10% O_2_ and increasing CO_2_ partial pressure. The dotted line indicates the terminal OD_600_ of WT *C. necator* at 48 hr in 1.5% CO_2_. Abbreviations: Hn = *Halothiobacillus neopolitans* Ba = *Bacillus anthracis* Vc = *Vibrio cholera* Fm = *Ferrovum myxofaciens* Af = *Acidothiobacillus ferrodoxans* SbtA from *Synechococcus elongatus* PCC 6301, BicA from *Synechococcus* sp. PCC 7002. **Panel B)** Representative data using the 1.5% CO_2_ partial pressure data from Panel A.

### The hnDAB2-expressing strain grows similarly to WT in constant-flow bioreactors but underperforms at high CO_2_

We next studied the growth profiles of our strains in constant-flow gas bioreactors under three conditions to assess the industrial relevance of our findings. The hnDAB2/ΔCA strain performed similarly to WT cells in ambient air supplemented with 5% H_2_, while the ΔCA strain did not grow (**Figure 5A**). We also tested two bioreactor conditions with CO_2_ partial pressures that are saturating for rubisco (5% and 10% CO_2_ **Figure 5B** and **5C**, respectively). We found that the WT strain grew much better in elevated CO_2_ than the hnDAB2/ΔCA strain.

**Figure 5.**
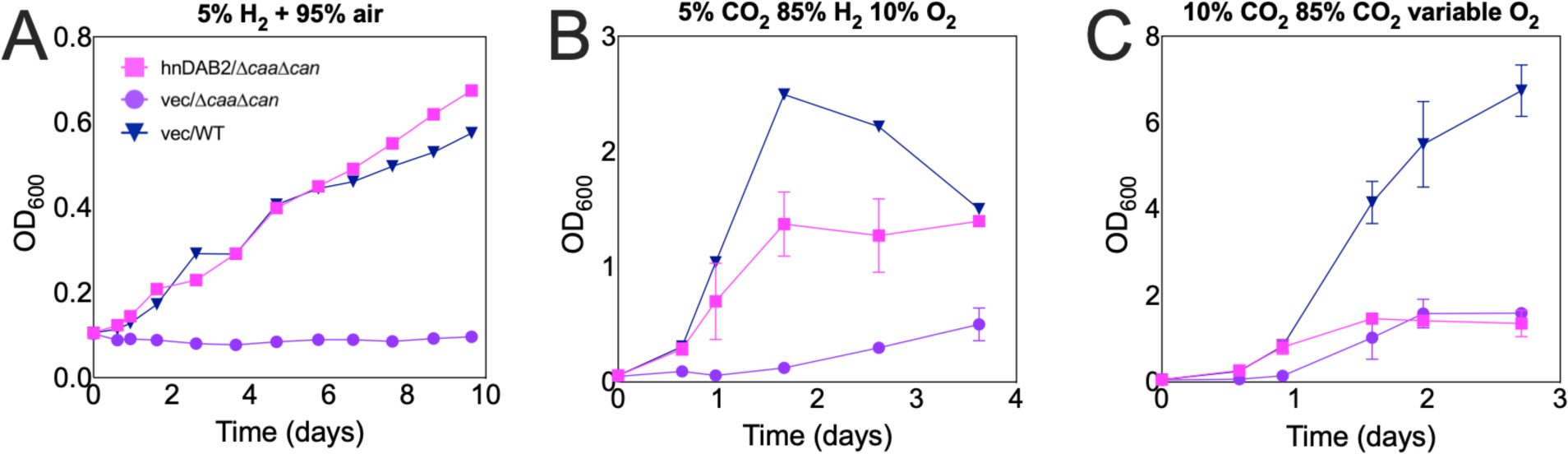
hnDAB2 restores growth of CA- strains to WT levels in ambient air in a continuously fed gas bioreactor supplemented with 5% H_2_ but underperforms WT in 5% and 10% CO_2_. The oxygen flow in bioreactors in panel B was gradually incremented from an initial value of 3% over the total gas flow to maintain DO levels of approximately >5%. Note the differences in scale of the y-axes when comparing panels.

### Chromosomal expression of hnDAB2 and heterologous rubisco expression

A plasmid-free system for expressing hnDAB2 would allow us to investigate the influence of heterologous rubisco expression in the context of DIC transport. To this end, the native genes for rubisco were deleted on chromosome 2, and pHG1 megaplasmid in the *ΔcaaΔcan* strain to produce a strain with genotype *ΔcaaΔcanΔcbbLS2ΔcbbLSp,* and hnDAB2 was added to the chromosome by a modified version of the Chassis-independent Recombinase-Assisted Genome Engineering (CRAGE) technique^44^. Briefly, a mariner transposon delivered a “landing pad” to the chromosome using a kanamycin selection. PacBio sequencing revealed that the landing pad landed in a 16S rRNA sequence, though no growth defects were observed in these strains. Next, a construct containing an expression cassette for hnDAB2 was delivered to the strain, and recombinants were selected for growth in ambient air (see supplementary information for details). The resulting strain (*ΔcaaΔcanΔcbbLS2ΔcbbLSp LP3::hndab2*) was used to assess the ability of rubisco homologs to recover autotrophic growth when paired with hnDAB2. We focused on bacterial rubisco homologs, as their requirements for assembly are well understood in *C. necator,* and the kinetic profiles of tested rubisco homologs are described in **Table 3**. All rubisco homologs were cloned onto a pBBR1-MCS plasmid driven by a native *C. necator* cbb_R_ promoter. We found that the rubiscos from *R. rubrum* and *R. sphaeroides* worked best in our system, though the fastest known bacterial rubisco from *Gallionella* sp. failed to grow^13^ (**Figure 6**). Expression of rubisco from *Synechococcus elongatus* PCC6301 did not rescue growth in *C. necator*, in agreement with previous findings^25^.

**Figure 6.**
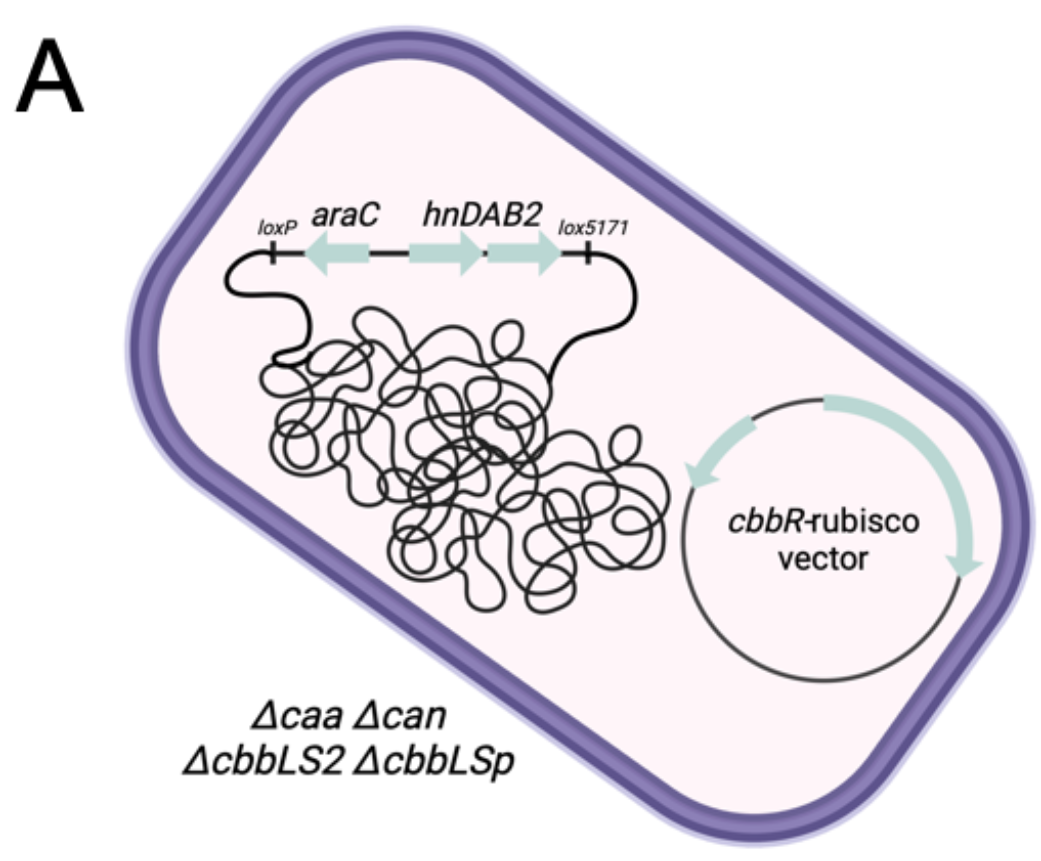
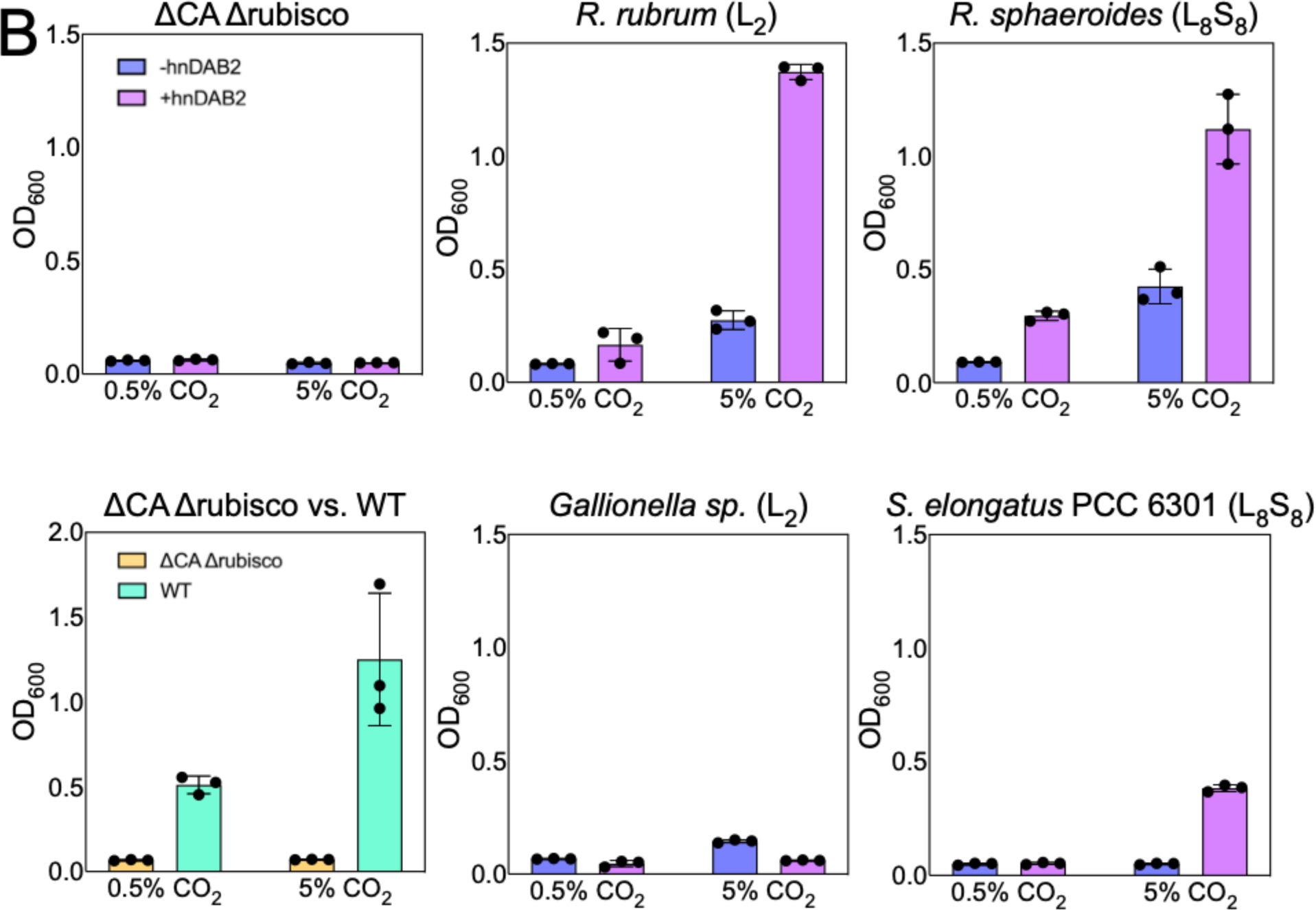
hnDAB2 activity improves growth in strains with heterologous rubisco expression. Panel A) Schematic describing the genetic configuration of strains used in these experiments. Strains either had a landing pad harboring a kanamycin resistance cassette or hnDAB2 under an arabinose promoter integrated into the *rpsL* locus. **Panel B)** Terminal endpoints for autotrophic growth at 48 hr in strains heterologously expressing various rubisco homologs. Cells were inoculated at an OD_600_ of 0.10 from heterotrophic precultures in 62% H_2_/10% O_2_ and indicated CO_2_. The source of the heterologous rubisco is indicated above each plot. Parentheses indicate the configuration of each rubisco in terms of large subunit (L) and small subunit (S) stoichiometry.

**Table 3.** Kinetic parameters of investigated rubisco homologs.

| Organism | Form | Type | $k_{cat}$ (s<sup>-1</sup>) | $K_M$ (μM) | $k_{cat}/K_M$ (s<sup>-1</sup>mM<sup>-1</sup>) | $S_{C/O}$ | Reference |
| --- | --- | --- | --- | --- | --- | --- | --- |
| *C. necator* H16 | IC | L<sub>8</sub>S<sub>8</sub> | 3.84 +/- 0.54 | 37 +/- 4 | 104 | 75 | 25,27 |
| *R. rubrum* | II | L<sub>2</sub> | 6.6 +/- 0.3 | 109 +/- 2 | 60 | 12.5 | 13 |
| *R. sphaeroides* | IC | L<sub>8</sub>S<sub>8</sub> | 6.7 +/- ND | 36 +/- ND | 186 | 62 | 13 |
| *S. elongatus* PCC 6301 | II | L<sub>8</sub>S<sub>8</sub> | 11.4 +/- 0.6 | 273 +/- 10 | 42 | 43.9 | 45 |
| *Gallionella* sp. | II | L<sub>2</sub> | 22.2 +/- 1.1 | 276 +/- 6 | 80 | 10 | 13 |

## Discussion

### Can is sufficient for autotrophic growth C. necator

We have demonstrated that Can is the primary CA allowing for the growth of *C. necator* H16 under heterotrophic and autotrophic conditions at low concentrations of CO_2_. Our conclusion agrees with multiple lines of published research^11,14^. The Caa enzyme has been proposed as a second CA involved in DIC metabolism in *C. necator*; however, we found only minor effects on growth in *caa* deletion mutants^24^.

### DIC transport suppresses bicarbonate starvation in Δcan strains

Our experiments demonstrate that several mechanistically distinct DIC transporters can complement strains of *C. necator* that are defective for the native CA Can. Our findings expand on discussions regarding bicarbonate/CO_2_ colimitation for autotrophic growth^39^. CO_2_ readily diffuses in and out of the cell (permeability coefficient P_C_ ≈ 0.1-1 cm/s)^30,46,47^. Consequently, the internal CO_2_ concentration is determined almost exclusively by the environmental conditions. At the lowest CO_2_ conditions tested (0.05%), we expect autotrophic growth to be limited by rubisco flux since the *C. necator* rubisco has a *K_M_*_(CO2)_ of 66 uM (in the presence of ambient air), corresponding to a CO_2_ gas concentration by volume of 0.16% in the headspace, as calculated by Henry’s law^48^. The rubisco will be saturated in the 0.5% CO_2_ condition and higher and should not be a growth limitation. In contrast to CO_2_, bicarbonate is a charged species that does not readily cross the cellular membrane. In the absence of CAs or DIC transport activity, the cellular flux of bicarbonate via the spontaneous hydration of CO_2_ is insufficient to meet the metabolic demands of anaplerosis. Therefore, we attribute the growth defect of *Δcan* to bicarbonate starvation and the subsequent rescue by DAB2, SbtA, and BicA to alleviating the bicarbonate bottleneck.

This finding is similar to a recent finding in C_3_ plants, where a CA defective mutant was fully capable of carrying out photosynthesis and only showed minor effects on the overall growth of the plant^40^. Our result regarding the importance of DIC accumulation in autotrophic metabolism is also supported by experiments showing that the expression of BicA and SbtA are well-suited to promote high cell density in high CO_2_ partial pressures (1.5-5% CO_2_). To our knowledge, this is the first report of functional heterologous expression of BicA, as several BicA homologs failed to rescue the growth of a ΔCA strain of *Escherichia coli*^49^.

### Completely rewired CO_2_ and HCO_3_^-^ metabolism

We have shown that the CO_2_/HCO_3_^-^ circuit in *C. necator* can be manipulated by replacing the native CAs and rubiscos with heterologous systems, including the expression of three different types of bicarbonate transporters and the functional expression of several types of rubisco homologs. Given that the examined rubiscos were chosen based on exemplary kinetic profiles, it is perhaps unsurprising that the expression of hnDAB2 largely governed the ability of *C. necator* to grow in autotrophic conditions. We found that *C. necator* can assemble functional rubiscos from several subtypes and geometries, which could pave the path toward highly efficient autotrophic bioproduction strains. *C. necator* features a form IC rubisco with L_8_S_8_ geometry, though the best-performing heterologous rubisco was the *R. rubrum* rubisco, which is form II and has an L2 geometry. The rubisco from R. rubrum is phylogenetically distinct from the rubisco in *C. necator* and has a broadly different kinetic profile (Table 3)^13,25^. Surprisingly, a rubisco variant from *Gallionella* sp. did not function in our strain despite having a similar kinetic profile to *R. rubrum* rubisco^13^. It is plausible that this variant’s expression is weak or the protein is not correctly folded in *C. necator*.

### Strains expressing hnDAB2 underperform in bioreactors with high concentrations of CO_2_

While strains expressing hnDAB2 outperformed WT in batch cultures at 1.5-5% CO_2_, we saw the opposite trend in constant-flow bioreactors. Given the robustness of the batch culture results, the bioreactor results were initially perplexing to us. The differences in equilibria between CO_2_ and HCO_3_^-^ in these two systems should be considered. In batch culture, we would expect that the majority of inorganic carbon is in the form of HCO_3_^-^ because mass transfer can occur over a prolonged period of time, and the equilibrium heavily favors HCO_3_^-^ formation. In contrast, the bioreactor flowed gaseous CO_2_ at two VVMs per minute, effectively giving a much higher concentration of gaseous CO_2_ compared to batch culture conditions. DAB2 is thought to work by hydrating CO_2_ on the cytoplasmic face of the complex by utilizing protons from the proton motive force (PMF). Given this mechanism, hnDAB2 may consume a large portion of the PMF in the higher concentrations of CO_2_ in the bioreactor, imparting a growth disadvantage relative to WT cells expressing Can CA, which produces bicarbonate from CO_2_ without consuming significant cellular resources such as PMF, ATP, or other cofactors. The consumption of PMF may explain why DABs are typically only found in organisms that grow in low pH, where protons are abundant, and CO_2_ is the dominant species^35^.

### Synthetic CO_2_ metabolisms and heterologous CCM expression: progress toward increased crop yield

Recently, there has been interest in optimizing carbon fixation pathways in organisms by expressing heterologous rubiscos or introducing CCMs, especially in crops, to increase food production. Our work is an important stepping stone for expressing a full CCM in *C. necator,* and we propose that our system is ideal for rapidly prototyping synthetic CO_2_ fixation systems and CCMs for further engineering in plants^50–52^. In contrast, eukaryotic hosts and many cyanobacteria are difficult and time-consuming to engineer. On the other hand, common bacterial hosts such as *E. coli* are not appropriate for screening for autotrophic growth advantages^53–55^. Using an *E. coli* strain dependent on the detoxification of RuBP to screen active rubiscos often gives false positives^28^. The work here and in ref. (25) show the utility of using *C. necator* H16 for high-throughput *in vivo* screening of rubisco performance, as the current methodology for biochemical screening of rubisco performance *in vitro* is labor and time-intensive.

## Conclusion

This work highlights the dual role that CO_2_ and its conjugate base, HCO_3_^-^, have in *C. necator*’s metabolic network. We have demonstrated extensive flexibility in *C. necator* autotrophic metabolism by the heterologous expression of many different types of bicarbonate acquisition systems and rubisco homologs. We showed that the predominant CA required for autotrophic growth is Can, which many DIC transporters can replace. Our findings support the notion that the primary role of Can during autotrophic growth is for bicarbonate accumulation, to be used as a cofactor for many carboxylases required for central metabolism. We propose that *C. necator* is an ideal system for engineering novel and synthetic CO_2_ metabolisms, such as PEP-driven carboxylation cycles or other pathways that are yet to be realized^56,57^.

## Methods

### Genetic manipulation and strain storage

Cloning fragments were amplified with Phusion® High-Fidelity DNA Polymerase (NEB, Ipswich, MA, USA) or Q5® High-Fidelity DNA Polymerase (NEB, Ipswich, MA, USA). PCR products were digested with Dpn1 (NEB, Ipswich, MA, USA) to destroy the original template and were purified using the QIAquick Gel Extraction Kit (Qiagen, Hilden, Germany). Ligation and assembly of DNA fragments were performed with NEBuilder® HiFi DNA Assembly Master Mix (NEB, Ipswich, MA, USA) according to the manufacturer’s protocol. The transformation was carried out via heat shock (CaCl_2_ method) into *E. coli* S17-1 cells. Cloned plasmids were purified using the QIAprep Spin Miniprep Kit (Qiagen, Hilden, Germany). Whole plasmid sequencing was performed by Primordium Labs (Arcadia, CA).

*C. necator H16* was routinely grown in LB media for cultivation and genetic procedures. Genomic deletions were made using a Kanamycin-SacB counterselection scheme using the suicide vector pKD18, and were performed by conjugation between plasmid-carrying *E. coli* S-17 strains and selected *C. necator H16* strains on LB agar plates and incubated overnight at 30°C. Conjugate cells were isolated on LB agar plates in the presence of 300ng*mL^-1^ kanamycin and 40ng*mL^-1^ gentamicin and incubated until *C. necator* colonies were observed. To create HCR strains, such as *Δcan*, plates were incubated in plastic gas bags containing saturating CO_2_ at 30°C. For strain storage, cells were inoculated into LB medium with appropriate antibiotics and were incubated overnight. Cells were pelleted to concentrate the biomass and were resuspended in 15% (v/v) glycerol, placed in cryogenic tubes, and stored at -80°C.

### Chassis-independent Recombinase Assisted Genome Engineering (CRAGE) method for C. necator H16

The CRAGE method was carried out as previously described, with modifications described in the SI^44^. Briefly, the plasmid encoding the mariner-based transposon harboring the landing pad (pW17) was modified to remove Cre recombinase and T7 RNA polymerase from plasmid pW17’’_noCreReg, and selection of the altered landing pad was carried out by conjugation into strain JP2281 (*ΔcaaΔcanΔcbbLS1ΔcbbLSp*), followed by selecting for growth on 200 µg/mL kanamycin in saturating CO_2_. PacBio sequencing was carried out at the Joint Genome Institute in Berkeley, CA, to identify the landing pad location in 16S rRNA ( 5’ CCAGCTACTGATCGTCGCCTTGGTAGGCTTTTACCCCACCAACTA::landing_pad:: TAGCTAATCAGACATCGGCCGCCCTGTAGCGCGAGGCCTTGC 3’ ). DAB2 was assembled into pW5Y under the *araBAD* promoter using yeast recombination cloning and was sequenced by the DIVA team at the Agile BioFoundry^58,59^. pW5Y.DAB2 was then conjugated into strain (JP2415 *C. necator ΔcaaΔcanΔcbbLS2ΔcbbLSp LP3*), and recombinants with DAB2 in the chromosome were selected for by growth on LB plates with no antibiotics in ambient air.

### Autotrophic growth in batch culture

*C. necator* strains were incubated overnight in 5 mL LB containing 200 µg/mL kanamycin (to retain plasmids) in 20 mL crimp-top tubes, supplemented with 10% CO_2_. Cultures were washed three times in Cupriavidus Minimal Media (CMM) and were inoculated into 150 mL serum vials in 5 mL of CMM without a carbon source. The vials were sealed and vacuumed. Gasses were transferred from gas sampling bags using syringes and needles with 62% H_2_ and 10% O_2_, and indicated partial pressures of CO_2_. Cells were grown at 30°C at 200 RPM for 48 hours and were then measured for terminal OD_600_ using a Molecular Devices® SpectraMax M2 spectrophotometer. The CMM contained 4.614 g/L Na_2_HPO_4_, 4.019 g/L NaH_2_PO_4_, 1.0 g/L NH_4_Cl, 0.455 g/L MgSO_4_*H_2_O, 0.453 g/L K_2_SO_4_, 0.047 g/L CaCl_2_, and 1 mL/L trace minerals solution (0.48 g/L CuSO_4_*5H_2_O, 2.4 g/L ZnSO_4_*7H_2_O, 2.4 g/L MnSO_4_*H_2_O, 15 g/L FeSO_4_*7H_2_O). The media was supplemented with 100 µg/mL kanamycin for experiments requiring the retention of pbadt-derived plasmids. Plasmids were not induced with L-arabinose for any of the experiments reported here, as we found basal expression rates of DIC transporters and CAs to be sufficient for function using pbadt vectors. However, L-arabinose (0.02%) was added to cultures to express hnDAB2 from the chromosome in rubisco swapping experiments at the initiation of batch autotrophic growth.

### Autotrophic growth in gas bioreactors

Cells were pre-cultured autotrophically in batch with 20 mL CMM containing 100 µg/mL kanamycin with 130 mL headspace containing 62% H_2_, 10% O_2_, and 10% CO_2_ balanced with N_2_, grown at 30°C with 200 RPM shaking for 48 hours. Growth of strains in bioreactors was performed using a bioXplorer® 400P system (HEL Ltd, UK) and WinIso® software for online monitoring and control. Each bioreactor was equipped with pH, dissolved oxygen (DO), temperature, and pressure controllers. The reactors were sterilized in an autoclave (120°C, 30 min), and filtered-sterilized autotrophic medium with kanamycin (100 μg mL^-1^ ) was batched into each. Temperature was maintained at 30°C and agitation at 1000rpm. The initial OD_600_ for all bioreactor experiments was between 0.05 and 0.10.

Experiments for growth in ambient air (i.e. Figure 5A) were performed with H_2_ (5%) (99.999% purity, Linde, US) and air (95%) continuously fed employing gas mass flow controllers and a total gas flow of 150 mL*min^-1^. The total working volume of the reactors was 300 mL.

Experiments using high CO_2_ levels were conducted with either 5% CO_2_ with 10% O_2_ and 85% H_2_ (i.e. Figure 5B), or 10% CO_2_ and variable O_2_, with a starting flow of 3.5% O_2_ and 86.5% H_2_ (i.e. Figure 5C). The total flow was 200 mL*min^-1^, and the reactor’s net volume was 250 mL. Oxygen flow was adjusted as consumed to maintain dissolved oxygen saturation levels under 30% to minimize any possible inhibition of autotrophic growth. Growth was measured by regularly sampling 2 mL of medium and analyzing optical density with a spectrophotometer in 10 mm pathlength cuvettes.

### Proteomics analysis

*C. necator* autotrophic cultures were collected at 48 hours of growth at 30°C and 200 RPM in 150 mL seared serum bottles containing 5 mL of microbial culture in 62% H_2_, 10% O_2_, and indicated concentrations of CO_2_. Cells were harvested and stored at -80 °C until further processing. Protein was extracted from cell pellets, and tryptic peptides were prepared by following the established proteomic sample preparation protocol (Chen et al.,. Briefly, cell pellets were resuspended in Qiagen P2 Lysis Buffer (Qiagen, Germany) to promote cell lysis. Proteins were precipitated with the addition of 1 mM NaCl and 4 x vol acetone, followed by two additional washes with 80% acetone in water. The recovered protein pellet was homogenized by pipetting mixing with 100 mM ammonium bicarbonate in 20% methanol. Protein concentration was determined by the DC protein assay (BioRad, USA). Protein reduction was accomplished using 5 mM tris 2-(carboxyethyl)phosphine (TCEP) for 30 min at room temperature, and alkylation was performed with 10 mM iodoacetamide (IAM; final concentration) for 30 min at room temperature in the dark. Overnight digestion with trypsin was accomplished with a 1:50 trypsin:total protein ratio. The resulting peptide samples were analyzed on an Agilent 1290 UHPLC system coupled to a Thermo Scientific Orbitrap Exploris 480 mass spectrometer for discovery proteomics^60^. Briefly, peptide samples were loaded onto an Ascentis® ES-C18 Column (Sigma–Aldrich, USA) and were eluted from the column by using a 10-minute gradient from 98% solvent A (0.1 % FA in H2O) and 2% solvent B (0.1% FA in ACN) to 65% solvent A and 35% solvent B. Eluting peptides were introduced to the mass spectrometer operating in positive-ion mode and were measured in data-independent acquisition (DIA) mode with a duty cycle of 3 survey scans from m/z 380 to m/z 985 and 45 MS2 scans with precursor isolation width of 13.5 m/z to cover the mass range. DIA raw data files were analyzed by an integrated software suite DIA-NN^61^. The databases used in the DIA-NN search (library-free mode) are *E. coli* and *C. necator* latest Uniprot proteome FASTA sequences plus the protein sequences of the heterologous proteins and common proteomic contaminants. DIA-NN determines mass tolerances automatically based on first-pass analysis of the samples with automated determination of optimal mass accuracies. The retention time extraction window was determined individually for all MS runs analyzed via the automated optimization procedure implemented in DIA-NN. Protein inference was enabled, and the quantification strategy was set to Robust LC = High Accuracy. Output main DIA-NN reports were filtered with a global FDR = 0.01 on both the precursor level and protein group level. The Top3 method, which is the average MS signal response of the three most intense tryptic peptides of each identified protein, was used to plot the quantity of the targeted proteins in the samples^62,63^.

### Data availability

The generated mass spectrometry proteomics data have been deposited to the ProteomeXchange Consortium via the PRIDE partner repository with the dataset identifier PXD051976^64^. Raw proteomics data is also available in SIproteomicsdata.xlsx.

Data collected during bioreactor runs is available in SIbioreactordata.xlsx

## Supporting information

10%co2_bioreactor_data

5%co2_bioreactor_data

Ambient_bioreactor_run

Top3_full_list_proteins

## Acknowledgments

We thank Luke Oltrogge for his extensive and thoughtful comments on the manuscript and Eric Sundstrom for his mentorship regarding the gas bioreactor experiments.

## Author Contributions

JP, experiment design, experiment execution, strain engineering, manuscript preparation

ET, experiment execution, strain engineering

STS, experiment execution

BF, experiment execution, strain engineering

ED, strain engineering

YC, proteomics

CJP, proteomics

DFS, funding, manuscript reviewing

SWS, funding, manuscript preparation

## Supplementary information

*Raw data:*

Top3_full_list_proteins.csv - Proteomics data CO2 step-up WT and hnDAB2/ΔCA 5%co2_bioreactor_data.xlsx - Bioreactor readouts for 5% CO2 run 10%co2_bioreactor_data.xlsx - Bioreactor readouts for 10% CO2 run Ambient_bioreactor_run.xlsx – Bioreactor readouts for 95% air 5% H2 run

*Strain identifiers for raw data:*

2071 – hnDAB2/Δ*caaΔcan*

2073 – vector/*ΔcaaΔcan*

2074 – vector/WT

## Modification of CRAGE technique

The CRAGE technique was modified because C. *necator* proved to be a challenging organism to engineer using this method. The transposon carrying the landing pad (expressing Cre recombinase) was delivered to strain JP2281 by conjugation and was selected for 200 µg/mL kanamycin. However, the *loxP* site was destroyed in 6/6 isolates, while the *lox5171* site remained intact (**Figure S1**). Removal of Cre recombinase from the landing pad resolved this issue (pW17’noCre). Instead, we expressed Cre transiently on a non-replicating delivery vector that encoded the hnDAB2 expression cassette (pW5Y-dab2). Further, we found that apramycin and chloramphenicol counterselection schemes failed in *C. necator* for CRAGE selection. We opted to use hnDAB2 or Can as a selection in the context of ambient CO2 growth, which resulted in recombinant hnDAB2+ cells.

## Complementation in a quadruple CA knockout

To assess whether Can operates alone or in combination with other CAs, we deleted all four carbonic anhydrases (ΔΔΔΔ) and assessed autotrophic growth of strains expressing Can or hnDAB2.

## Growth on HCO3- as a sole carbon addition

Similar growth trends were observed when HCO_3_^-^ was added as the sole carbon source, though this is unsurprising considering that HCO_3_^-^ equilibrates with CO_2_ rather quickly (minutes to hours). We added bicarbonate in equimolar concentrations to CO_2_ in previous experiments and observed very similar growth patterns to previous experiments. While the ΔCA strain did not grow, all strains complemented with a DIC transporter grew similarly using KHCO_3_ or equimolar CO_2_ (**Figure S3**). However, stains did not grow well at 61 mM KHCO_3,_ likely caused by osmotic pressure imparted by excessive K^+^, as the addition of 61 mM KHCO_3_ nearly doubled the salt concentration of the media. Similarly, excess K^+^ may negatively impact the performance of bicarbonate transporters, particularly in the case of BicA and SbtA, which are Na^+^ transporters and may be competitively inhibited in the presence of excessive K^+^.

**Figure S1.**
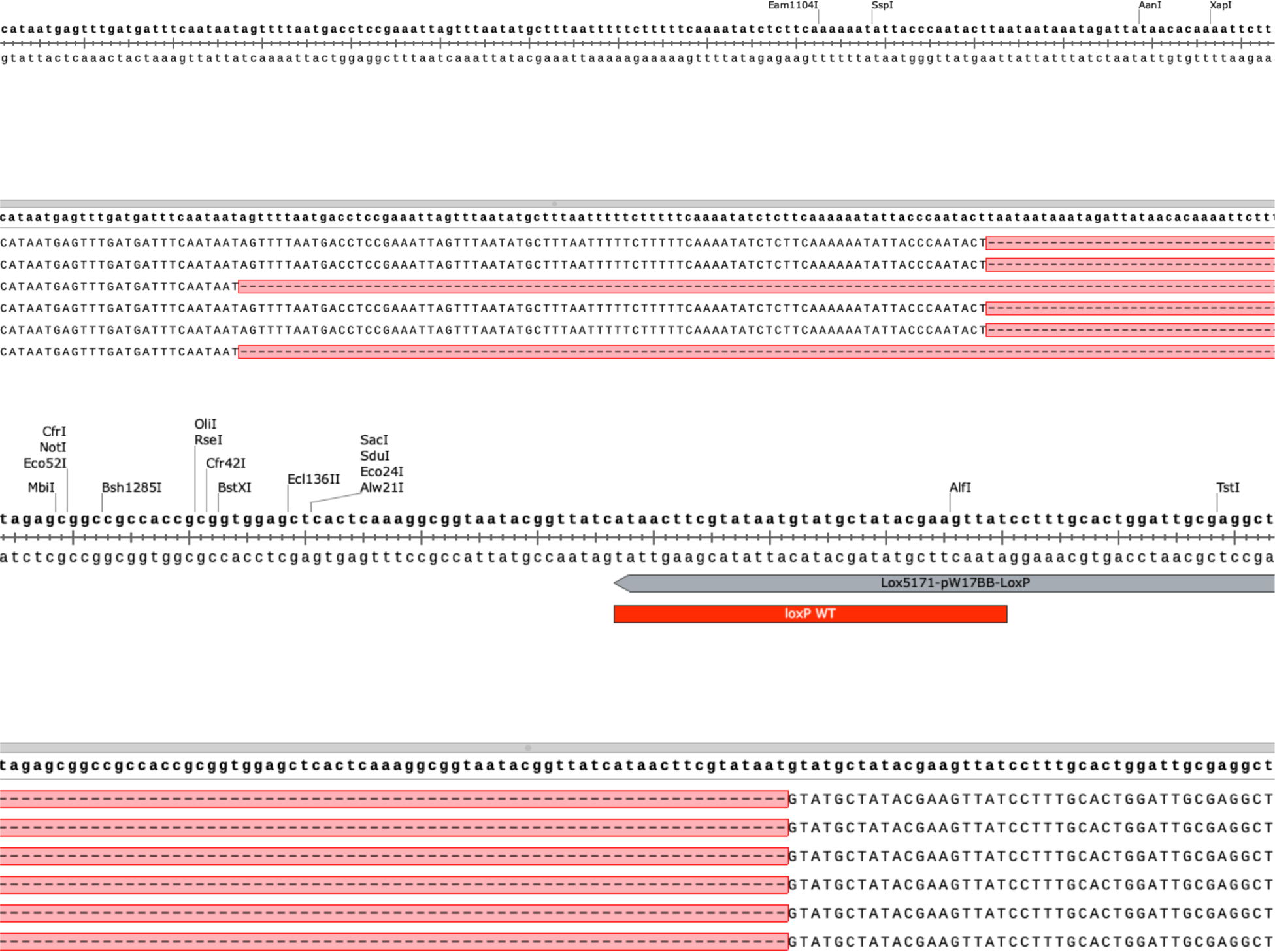
Truncations were observed in the loxP site when delivering a full-length CRAGE landing pad that expresses Cre recombinase. The landing pad for CRAGE was disturbed at the critical *loxP* site in 6/6 isolates. Four isolates contained a 210 bp deletion, creating a ‘TCAAAAAATATTACCCAATACT **GTATGCTATACGAAGTTAT**‘ scar. Two of the six isolates contained a 294 bp deletion, creating a ‘TAAATCCATAATGAGTTTGATGATTTCAATAAT **GTATGCTATACGAAGTTAT‘** scar.

**Figure S2.**
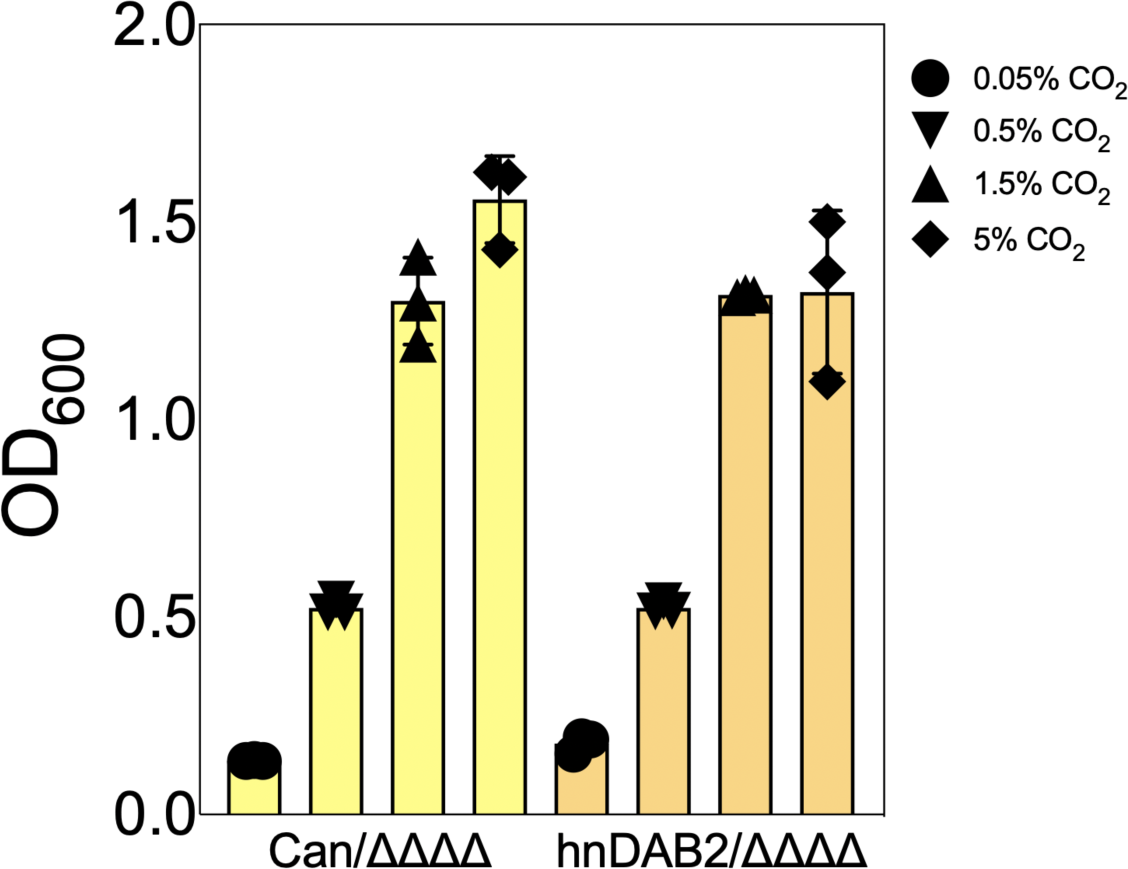
**hnDAB2 is able to complement the quadruple CA knockout mutant and allows for similar growth to the pCan complemented strain.**

**Figure S3.**
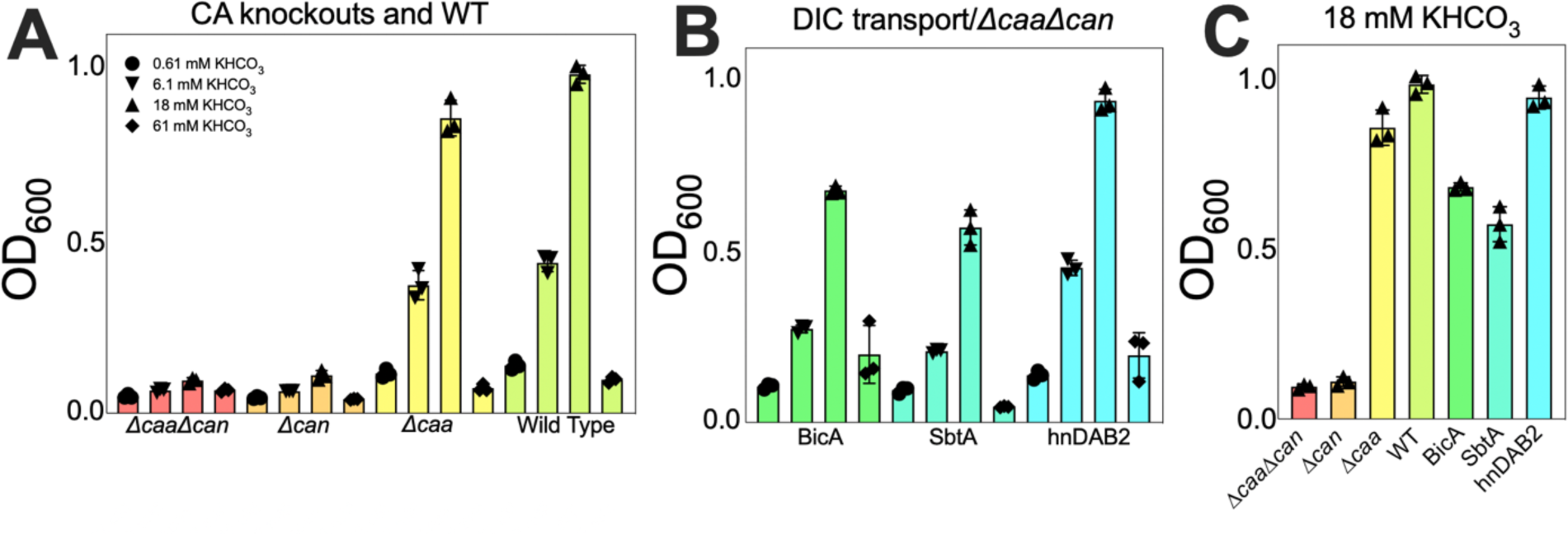
Strains with Can CA or a DIC transporter but no Can CA grow with HCO3^-^ as the only carbon addition to the media. Showing data for endpoints after 48 hours of growth of autotrophic growth in 62% H, 10% O2, and indicated concentrations of KHCO3. **Panel A)** Strains that have *can* CA intact can grow with only bicarbonate in the media. **Panel B)** All three types of DIC transporters can complement for Can on HCO3^-^. **Panel C)** A comparison of the endpoints for all strains tested in 18 mM KHCO3.

